# IFNBoost: An interpretable computational model for identifying IFNγ inducing peptides

**DOI:** 10.1101/2025.01.21.634172

**Authors:** Inam ul haq azad, Muhammad Saqib Sohail, Ahmed Abdul Quadeer

## Abstract

**Motivation:** Interferon-gamma (IFN*γ*) is a pivotal cytokine that coordinates various aspects of the immune response, notably enhancing T-cell activation, clearing intracellular pathogens, and providing long-term immune protection. Identification of IFN*γ*-inducing peptides is essential for the advancement of peptide-based vaccines and immunotherapies; however, the experimental determination of these peptides is hampered by the large number of potential peptide candidates present in pathogen proteins.

**Results:** In this study, we present IFNBoost, a machine learning model developed to accurately predict IFN*γ*-inducing peptides by leveraging existing immunological datasets, including both peptide sequences and associated metadata. IFNBoost demonstrates impressive performance metrics, achieving an accuracy of 0.819, an F1 score of 0.798, and a Matthew’s correlation coefficient (MCC) of 0.634. Evaluation against independent datasets demonstrates that IFNBoost surpasses all current models for predicting IFN*γ*-inducing peptides, highlighting generalizability of the model. Our comprehensive analysis indicates that, in addition to peptide sequences, metadata features such as the source organism and host significantly enhance predictive accuracy. The predictions produced by IFNBoost have the potential to guide rational vaccine design, thereby improving vaccine efficacy via precise identification of peptides that elicit the desired cytokine responses.

**Availability and implementation:** To improve the accessibility and utility of our model, we have developed a web application available at https://ifnboost.streamlit.app/.

## Introduction

Cytokines are essential signaling molecules that regulate the immune response by stimulating or inhibiting the activation, proliferation, and/or differentiation of various cells. Cytokines such as IL-2 and IFN*γ* promote the differentiation and activity of natural killer cells and cytotoxic T cells (CTLs), leading to a cytotoxic immune response [1]. In contrast, cytokines like IL-4 and IL-5 regulate B cell activity and promote antibody response. Their importance extends to vaccine development, wherein the rational design of vaccines underscores the need of identifying antigens capable of inducing specific cytokines that promote a specific immune response [2, 3]. This makes it essential to identify epitopes (antigenic regions) from pathogens that can stimulate the production of a particular cytokine.

Among the various cytokines, IFN*γ* plays a key role in both innate and adaptive immunity by activating macrophages, enhancing antigen presentation, and promoting the differentiation of T cells into T-helper 1 (Th1) cells [4]. It also has significant antiviral and antibacterial properties [5], and acts as a therapeutic and diagnostic tool in diseases like cancer and tuberculosis [6, 7]. For example, Figure 1 depicts the working of T cells upon recognition of an infected cell, where T-helper cells secrete IFN*γ* cytokine and promote CTL response, highlighting the significance of identifying IFN*γ* inducing peptides for the design of peptide-based vaccines or immunotherapies [8]. In designing a peptide-based vaccine for any virus, one of the main challenges is to determine peptides capable of promoting specific immune cells (antibodies or T cells) that are known to protect against the disease. Identifying such peptides requires comprehensive experimental investigation via different assays (e.g., ELISPOT, ICS, etc.), though this is prohibited by the huge number of possible peptides in a viral protein of interest.

**Figure 1:**
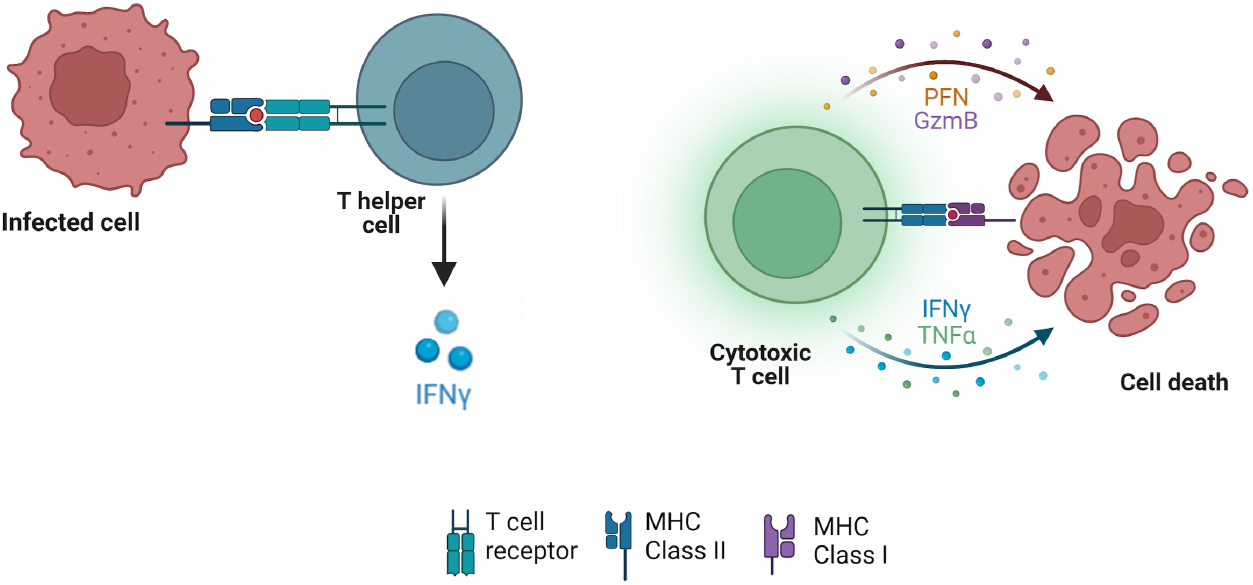
T cell function upon recognition of infected cells. When T helper cells identify an infected cell, they release IFN*γ* cytokine. This activates the CTLs, which induce apoptosis (programmed cell death) in the infected cells. CTLs release cytotoxins, including perforin (PFN) and granzymes (GzmB) to facilitate this process. Perforin forms pores in the target cell membrane, allowing granzymes to enter the target cell and trigger apoptosis. This figure was generated with BioRender.com.

To confront this challenge, computational methods are utilized to harness the available experimental data for different types of virus peptides and the immune cells they promote. In-silico tools like IFNepitope [8], ILeukin10Pred [9], Meta-IL4 [10] have been developed to predict the cytokine-inducing peptides. To our knowledge, IFNepitope [8] and IFNepitope2 [11] are the only existing tools for predicting IFN*γ* inducing peptides. These methods utilise only the epitope sequence information for making predictions.

However, these methods lack thorough and systematic evaluation of their predictive performance from a machine learning (ML) perspective. Moreover, their generalizability to independent datasets has not been rigorously tested. Key questions that remain unanswered include: Is the peptide sequence alone sufficient to distinguish IFN*γ* inducers and non-inducers? How important are epitope-specific metadata features like host and source organism for prediction? Addressing these uncertainties is critical to improving the reliability and applicability of these models.

In this work, we address these challenges by developing IFNBoost, a machine learning (ML) model for predicting IFN*γ* inducing peptides, trained and tested on experimental data from the Immune Epitope Database (IEDB) [12]. IFNBoost is distinct from existing models as it incorporates three metadata features of host, source organism species, and assay method as additional inputs, along with epitope sequence. It achieves excellent performance based on all evaluation metrics in distinguishing IFN*γ* inducing peptides from a given set of peptides. We also benchmark IFNBoost against existing tools and assess their performance and generalizability on an independent dataset. A detailed interpretation of IFNBoost is performed to understand the importance of different metadata features in accurate IFN*γ* response predictions. We provide a web server for IFNBoost that features a graphical user interface for text-based queries of epitopes. Overall, this work provides a systematic framework for the prediction of IFN*γ* inducing epitopes, contributing to the broader goal of designing effective vaccines.

## Materials and methods

### Dataset acquisition and preprocessing

We accessed IEDB to retrieve T cell epitope data on July 13, 2023. Since our interest was in IFN*γ* inducing MHC class II-restricted T cell epitopes, data was downloaded by selecting ‘T cell’ under the ‘Assay’ category and ‘Class II’ under ‘MHC restriction’, while leaving all other search parameters (Epitope, Epitope source, Host, Disease) set to ‘Any’.

The downloaded dataset comprised 2,25,023 epitopes obtained from diverse T cell assays and reported to secrete different types of cytokines. We identified 68,651 epitopes for which information regarding IFN*γ* cytokine release was available. From these epitopes, only those containing valid sequences composed of the standard 20 amino acids were retained. Different lengths of MHC Class II binders have been reported in the literature [13, 14, 15]. In the downloaded dataset, more than 99% of epitope sequences had length of 31 amino acids or less (Supplementary Figure 1). To cover the maximum number of epitope sequences, we included epitopes ranging from 12 to 30 amino acids in length for further processing. Additionally, we identified three different entries for Homo sapiens namely ‘Homo sapiens Caucasian’, ‘Homo sapiens (human)’, and ‘Homo sapiens’. As we did not consider ethnicity in our study, we consolidated all of these entries under the single designation of ‘Homo sapiens’ for subsequent analysis.

Next, we focused on specific information relevant to our work: epitope sequence, epitope-species (source organism species), host, assay-method, and assay-qualitative measurement. The assay-qualitative measurement represents the IFN*γ* response of an epitope. Epitopes reported with both positive and negative IFN*γ* responses were considered in the positive IFN*γ* epitope class, following [8].

### Feature encoding

The preprocessing steps resulted in a total of 41,467 samples, wherein each sample consists of five pieces of information: epitope sequence, source organism species, host, assay-method, and assay-qualitative measurement. The assay-qualitative measurement was used as a class label in this study. Positive IFN*γ* responses were mapped to ‘1’, while negative responses were mapped to ‘0’, ensuring a consistent representation of data for binary classification. The input to the model consists of the remaining four features. Since we restricted epitope sequences to a maximum length of 30 amino acids, each sequence was represented as a vector of 30 entries. This, together with the other three features of host, source organism species and assay method, serve as the model input, comprising 33 entries (Supplementary Figure 2). The processed dataset consisted of 18,464 samples in the positive class and 23,003 samples in the negative class. This dataset was partitioned into training and testing sets using a 80:20 train-test split ratio which resulted in 33,173 samples in the training and 8,294 samples in the test set.

To provide numerical inputs to the ML model, the input features were encoded in a numerical format. We applied zero-padding to standardize the epitope sequences to a uniform length of 30 amino acids. Ordinal encoding was employed to map the input information to numeric values, which were then input into the XGBoost model. We established a threshold criterion to map the metadata features of host and source organism species, wherein the top categories were selected such that they collectively represent 99% of the epitope sequences. In the training data, the top 43 out of 157 hosts and 121 out of 264 source organism species accounted for this 99% representation. The top 43 hosts were assigned integer values from 1 to 43, while all other hosts were assigned the integer 44, designating them as the ‘others’ category. A similar approach was applied to the source organism species feature. For samples with unknown source organism species, these were also designated to the ‘others’ category of the source organism species, which also allows the handling of missing entries. In the training dataset, there were only seven assay methods, which we encoded with integer values ranging from 1 to 7. Detailed metadata feature mapping is listed in Supplementary File 1. The test set samples were encoded based on the scheme developed for the training set. Any novel categories present in the test set were assigned to the ‘others’ category.

### Model selection and hyperparameter optimization

The considered epitope dataset consists of tabular data, organized in columns containing information related to each epitope. Multiple studies [16, 17, 18] have shown that decision tree-based models outperform deep learning methods on various tabular datasets. Among the decision tree-based models, XGBoost [19] stands out for its ease of optimization and is often recommended for practical applications involving tabular data [17]. Hence, we chose the XGBoost model to predict IFN*γ* inducing epitopes.

XGBoost is an implementation of gradient boosted decision trees. The framework uses a gradient boosting approach, where each new tree is built to correct the errors made by the existing ensemble of trees. To optimize the XGBoost model, we considered 10 hyperparameters: *n estimators* represents the number of trees used to build the XGBoost ensemble, *learning rate* determines the step size at each boosting iteration, and *max depth* controls the maximum number of layers or levels in a decision tree. *λ* and *α* are the coefficients of L2 and L1 regularization which aid in controlling overfitting. *γ* controls the minimum loss reduction required to make a further partition on a leaf node of the tree, with a larger value of gamma lowering the complexity of the model, and reducing overfitting. *min child weight* is used to control the minimum number of observations required to create a new partition during the tree-building process, where a higher value of *min child weight* makes the model more conservative. *Tree method* determines the tree construction algorithm, with *subsample* controlling the fraction of training instances to be randomly sampled for each tree-building iteration (used to introduce randomness in the model) and *colsample bytree* controlling the fraction of features (columns) to be randomly sampled for each tree during the tree building process. We optimized these hyperparameters for different search spaces (see Supplementary Table 1). For *tree method*, we considered two choices: exact and auto.

Optimization was conducted using five-fold cross-validation (CV) of training data, employing the open-source Python package Optuna [20]. Optuna uses a Tree-structured Parzen Estimators (TPE) algorithm [21] which is based on the Bayesian update framework. This approach combines random exploration within a defined search space and subsequently minimizes the objective function based on the acquired knowledge from previous iterations.

### Details of comparative approaches

We compared IFNBoost with existing state-of-the-art methods for predicting IFN*γ* inducing peptides. In 2013, [8] introduced a method called IFNepitope for this purpose. Recently, the same group released an updated version, IFNepitope2 [11]. Both methods are accessible via web servers and serve as benchmarks for evaluating the performance of our proposed IFNBoost method.

A fundamental difference between these methods is that they rely solely on epitope sequence information, whereas we incorporate metadata alongside epitope sequence encoding as inputs to our model. Their approach encodes epitope sequences based on di-peptide composition (DPC), which represents the frequency of all possible amino acid pairs. Predictions are made using a hybrid method that combines an ML model with a sequence similarity approach [8,12]. Notably, IFNepitope2 is designed to predict IFN*γ* inducing peptides specifically for two hosts, human and mouse, while IFNepitope is not restricted to specific host organisms. IFNepitope2 offers two separate web servers for predicting epitopes associated with mouse and human hosts. In addition, IFNepitope2 is limited to epitopes with lengths of up to 20 amino acids.

For model comparison, we downloaded T cell epitope data from IEDB that was uploaded/reported in 2024. This dataset serves as an independent test set, as all three evaluated methods (IFNBoost, IFNepitope, and IFNepitope2) were developed using IEDB data available up to 2023 or earlier. The downloaded data was preprocessed using the same criterion mentioned in Dataset acquisition and preprocessing section. We obtained 1,012 samples to report the performance of IFNBoost (see Results). To evaluate IFNepitope, we retained 944 unique epitope sequences from these samples. For IFNepitope2, we categorized the sequences by host species, testing 145 human-associated and 709 mouse-associated samples on their respective web servers. The outputs were merged to create an aggregated confusion matrix, combining the confusion matrices from both models. Using this aggregated matrix, we calculated and reported the overall performance of IFNepitope2 on the 2024 dataset across different evaluation metrics, as detailed in the Performance metrics section below.

### Testing IFNBoost by varying the input features

We conducted tests to analyze the importance of input features in predicting the IFN*γ* response of epitopes using IFNBoost. Initially, we evaluated the model’s performance using 30 input features derived exclusively from amino acid encoding, excluding meta-data. In this scenario, unique epitope sequences were treated as distinct samples, resulting in a dataset of 34,292 epitopes. Following an 80:20 train-test split, XGBoost was trained on 27,433 samples, with 6,859 reserved for testing. Next, we tested the impact of including individual metadata features alongside sequence encoding, leading to 31 input features per test case. And for completeness, we tested combinations of two metadata features with sequence encoding, resulting in 32 input features.

### Different amino acid encodings

In IFNBoost, we have used the ordinal encoding scheme to represent the epitope sequence. Ordinal encoding assigns numerical values from 1 to 20 to represent the 20 unique amino acids within sequences up to a length of 30. Zero padding is utilized to represent shorter amino acid sequences, denoted by the digit 0. Other encodings that we tested include:

1. Di-peptide composition (DPC): This encoding was used in developing IFNepitope and IFNepitope2 models, which represents the frequency of occurrence of all possible amino acid pairs (dipeptides) in a given protein sequence. This generates a vector of length 400 for each epitope sequence.
2. Physicochemical properties based encoding: This encoding scheme categorizes all amino acids into five classes: apolar, aromatic, polar neutral, acidic, or basic [22]. These classes are represented by numbers from 1 to 5, generating a vector of length 30 for each epitope sequence.
3. Betts and Russel classification based encoding: This encoding scheme is based on physical, chemical and structural properties of side chains [23]. It classifies amino acids into aliphatic, aromatic, neutral, negatively charged, positively charged, and small, which are represented by numbers from 1 to 6. Similar to the physicochemical properties based encoding, this encoding also generates a vector of length 30 for each epitope sequence.

### Performance metrics

IFNBoost and other methods were evaluated on a binary classification task to predict whether an epitope is an IFN*γ* inducer or non-inducer. We assessed the performance of these methods using five standard evaluation metrics: Sensitivity, Specificity, Accuracy, F1 Score, and Matthews Correlation Coefficient (MCC). Sensitivity and Specificity measure the model’s ability to correctly identify positive (IFN*γ* inducers) and negative instances (non-inducers), respectively. Accuracy provides an overall measure of correctness. The F1 score and MCC are particularly important metrics as they account for class imbalance within the dataset. These metrics were calculated using the confusion matrix, which encompasses true positives, false positives, true negatives, and false negatives.

## Results

### IFNBoost: A machine learning model for predicting IFN*γ* inducing peptides

Extreme Gradient Boosting (XGBoost) framework ([19]) was employed to develop IFNBoost, a predictive model for detecting IFN*γ* inducing epitopes. The input to this model is a vector of 33 features, where the first three features represent the host, source organism species, and assay method, while the remaining 30 features represent the amino acid encoding (Figure 2a). The dataset consists of 43,016 samples, with 44% classified as IFN*γ* inducing epitopes, reflecting a mild imbalance in the data distribution. Following an 80:20 train-test split of the entire dataset, we trained and optimized our IFNBoost model on 33,173 samples, and reported the performance on the remaining 8,294 (test) samples. IFNBoost achieved an area under the receiver operating characteristic (AUROC) curve of 0.898 and an accuracy of 0.819 on the test set (Figure 2b). With an area under the precision-recall curve (AUPRC) of 0.870 (Figure 2c), the model demonstrates strong predictive capability for IFN*γ* inducers. IFNBoost achieved sensitivity of 0.806 and specificity of 0.829, indicating the model’s ability to accurately identify both true positives (IFN*γ* inducers) and true negatives (non-inducers) respectively (Figure 2d). This highlights its balanced performance in distinguishing between the two classes. Additionally, the F1 score of 0.798 reflects a balanced trade-off between precision and recall, while the MCC of 0.634 indicates the effectiveness of the model in handling the dataset’s class imbalance (Figure 2d). Overall, the results suggest that IFNBoost can predict both the IFN*γ* inducing epitopes and non-inducing epitopes with comparable accuracy.

**Figure 2:**
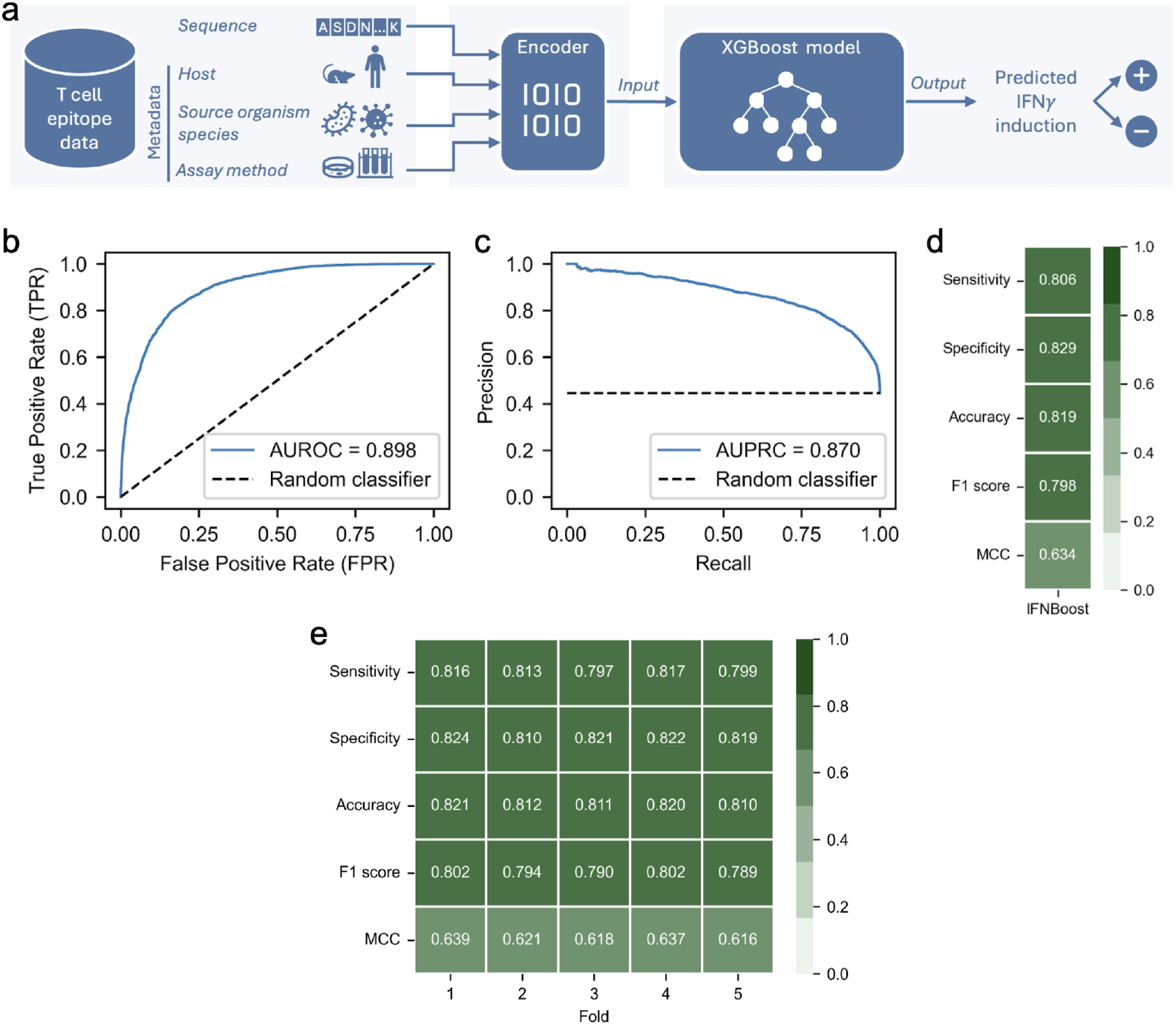
IFNBoost model for predicting IFN*γ* inducing peptides. (a) Overview of the IFNBoost model. (b-e) Performance evaluation of the IFNBoost model. (b) Receiver operating characteristic (ROC) curve illustrating the trade-off between TPR and FPR at various threshold settings. AUROC represents the area under the ROC curve and it ranges from 0 to 1. The diagonal line (random classifier) represents a baseline reference with an AUROC of 0.5. (c) Precision-Recall Curve (PRC) showing the trade-off between precision and recall across different classification thresholds. The random classifier’s precision is indicated at 0.44, corresponding to the proportion of positive samples in the dataset. (d) The heatmap represents the performance of the IFNBoost model on the test set. (e) Five-fold cross-validation results of IFNBoost. The complete training dataset, consisting of 33,173 samples was partitioned into 5 folds, and in each iteration, 4 folds were used for training, and 5th fold was left for testing. The model parameters were set the same as those used to report evaluation metrics in (b)-(d).

The epitope sequence dataset exhibits considerable diversity in terms of the meta-data features, encompassing 157 different hosts, 264 species of source organisms and 7 assay methods. However, the top 10 hosts and source organism species account for a significant proportion of the epitopes (Supplementary Figure 3), indicating that while the dataset is broad, certain features are over-represented. To test our model’s robustness and generalization ability across the dataset, we performed five-fold cross-validation (CV) using the complete training dataset of 33,173 samples. Simulations indicated that the five-fold CV results were consistent across all the folds (Figure 2e), demonstrating the model’s capacity to generalize effectively across epitope sequences with varying metadata features. We used the same optimized parameters of IFNBoost for each fold. To prevent data leakage between the training and test samples, feature encoding was learned from the training samples in each iteration and subsequently used to characterize the features of the test set. The model demonstrated low variation across the five folds, indicating consistent model performance.

### Generalizability of IFNBoost and benchmarking against existing models

IFNBoost was developed using the IEDB epitope dataset collected prior to 2024. For external validation of IFNBoost and to assess its generalizability on unseen epitope sequences (not present in the training data), we evaluated it against an independent dataset from IEDB which comprises T cell epitope data reported in 2024. IFNBoost demonstrated impressive performance on this independent dataset (Figure 3a), achieving an accuracy of 0.888. Remarkably, it maintained this high predictive performance with unseen epitope sequences, attaining an accuracy of 0.889.

**Figure 3:**
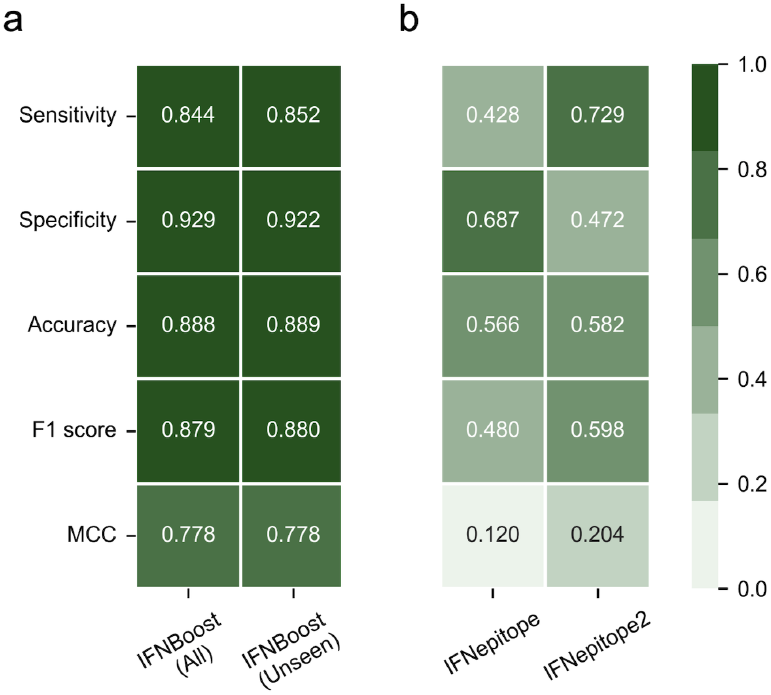
Generalizability of IFNBoost and comparison with other methods. (a) IFNBoost performance on the independent data from 2024. ‘All’ refers to 1012 samples obtained from the independent dataset after preprocessing, while ‘Unseen’ refers to 839 samples from the independent dataset not present in IFNBoost training. (b) Performance of existing methods on the independent data from 2024. IFNepitope was evaluated on 944 samples and IFNepitope2 on 854 samples (see **Materials and Methods** for details).

To benchmark IFNBoost, we compared it with the two existing methods that predict IFN*γ* inducing peptides, IFNepitope [8] and IFNepitope2 [11], on the same independent dataset. These two methods are based on sequence information only and do not incorporate the metadata. We utilized their respective web servers to make predictions on the independent dataset (see **Materials and Methods** for details). IFNBoost outperformed IFNepitope and IFNepitope2 across all evaluated performance metrics (Figure 3). It demonstrated higher sensitivity and specificity, indicating a greater ability to correctly identify both true positive and true negative instances of IFN*γ* inducers and non-IFN*γ* inducers, respectively. IFNBoost achieved a substantially high accuracy of 0.888 compared to 0.566 for IFNepitope and 0.582 for IFNepitope2, demonstrating its overall enhanced predictive performance. Additionally, IFNBoost surpassed the other two methods based on the F1 score and MCC. These results suggest that IFNBoost provides superior performance in identifying both IFN*γ* inducers and non-inducers.

### Interpretation of the IFNBoost model

Having established the strong performance of IFNBoost in predicting both IFN*γ* inducing and non-inducing peptides, we conducted a comprehensive analysis to evaluate the contribution of various epitope features to model predictions. Specifically, we assessed the significance of the epitope sequence, metadata features (host, source organism species, and assay method), epitope length, and different amino acid encoding schemes.

#### Importance of epitope metadata

Using only the epitope sequence (30 features) as input, the model achieved an accuracy of 0.653 (Figure 4a left panel). Adding each metadata feature progressively improved the model performance. The inclusion of the assay method alone improved the accuracy to 0.702, while the inclusion of the host alone led to an accuracy of 0.730. Incorporating the source organism species had the greatest effect on accuracy increase resulting in 0.756 accuracy. The inclusion of two metadata features in conjunction with epitope sequence further improved model performance compared to using individual metadata features. Specifically, the inclusion of host and source organism species achieved a relatively high accuracy of 0.809. Expanding the analysis to include all three metadata features resulted in the highest accuracy of 0.819, underscoring the critical importance of all three metadata features in IFNBoost for accurately predicting IFN*γ* inducing peptides.

**Figure 4:**
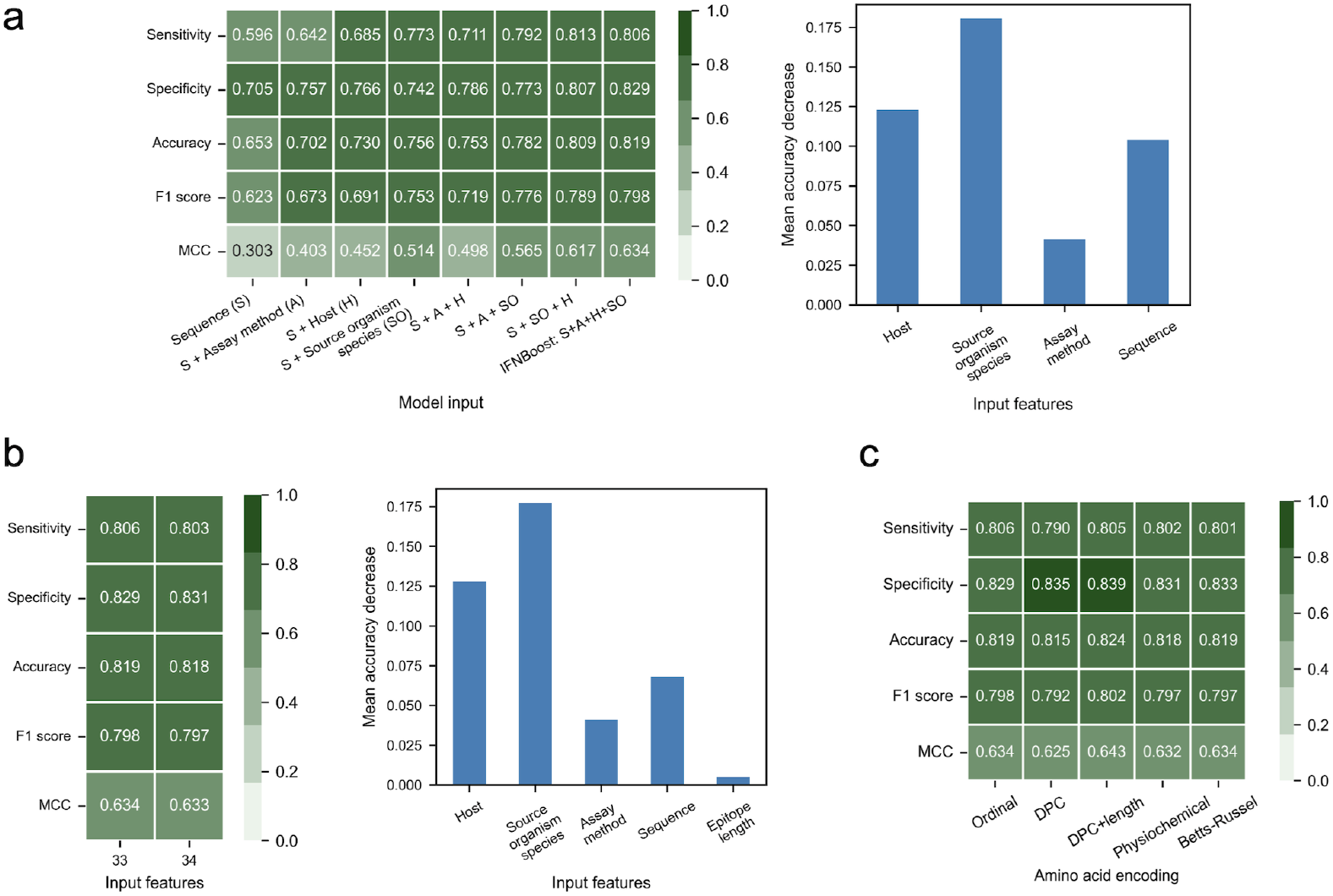
Interpretation of the IFNBoost model. (a) The importance of including epitope metadata on model’s performance. (Left panel) Performance comparison by varying the input features to the model. The inclusion of metadata features along with epitope sequence improves the classification performance of the model. (Right panel) Input feature importance of IFNBoost. The bar chart illustrates the permutation importance analysis of the input features for the model, conducted using scikit-learn ([24]) library’s ‘permutation importance’ function, considering 33 input features. ‘Sequence’ represents the aggregate scores of the 30 features of amino acid encoding. Each bar represents the mean accuracy decrease by randomly shuffling a single feature value. The analysis involved 10 repetitions, i.e., the permutation process is repeated 10 times for each feature. The importance of IFNBoost has been calculated on the same test set used to report results in Figure 2b-2d. (b) Impact of epitope length on model’s performance. (Left panel) Performance comparison of IFNBoost (33 input features) to the same model augmented with an additional input feature of epitope length (34 input features). (Right panel) Input feature importance of the model with ‘epitope length’ as an additional feature. The feature importance was calculated using the permutation importance method similar to subfigure a, considering 34 input features. (c) Model performance using different types of amino acid encodings. Ordinal encoding assigns numerical values to represent the 20 unique amino acids. DPC represents the di-peptide composition-based encoding of amino acids. DPC+length combines DPC with epitope length as an additional input feature. Physicochemical and Betts-Russel encodings group amino acids based on different properties (see **Materials and Methods** for details).

To complement this analysis, we assessed the importance of model input features for IFNBoost using the permutation importance analysis approach [25] (Figure 4a right panel). This analysis helps to identify which of the input features the trained model prioritizes in addressing the prediction problem. Consistent with the findings in Figure 5a, the source organism species emerged as the most critical feature, as indicated by the greatest mean accuracy decrease. The epitope sequence, consisting of 30 features, is more important than the assay method in predicting the IFN*γ* response. While individual features within the epitope sequence showed varying importance (Supplementary Figure 4), with positions 16 and 18 being particularly significant, their collective contribution is significant. Together, all 30 features provide essential sequence-level information that enhances the model’s predictive ability, highlighting the importance of integrating epitope sequence with metadata features.

**Figure 5:**
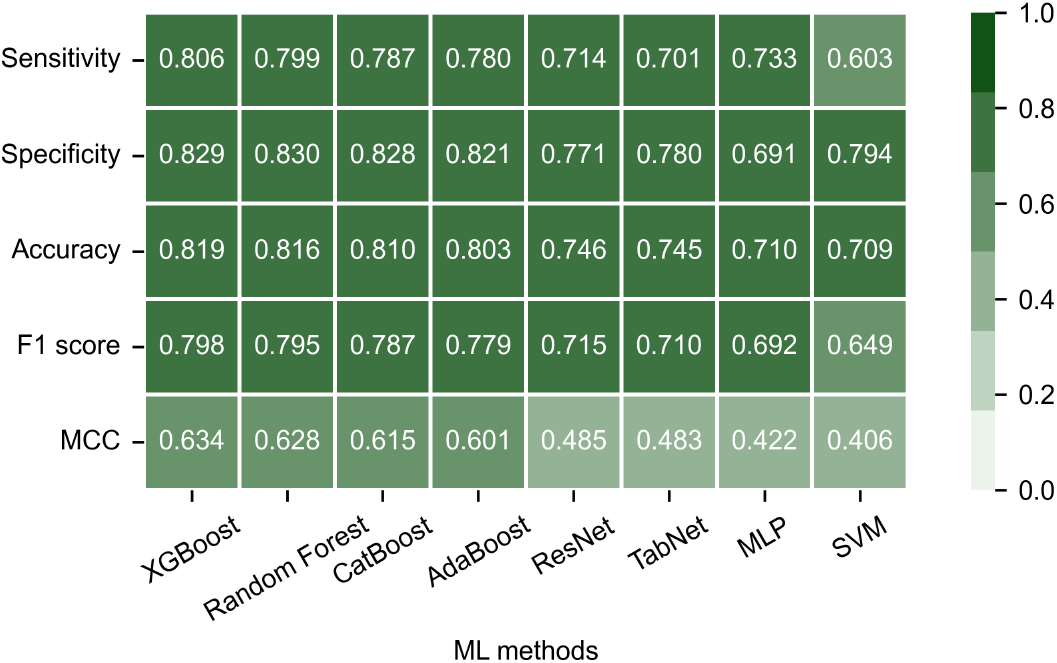
Performance comparison with other ML methods. The methods have been arranged in decreasing order of their performance. Random Forest, AdaBoost and SVM have been implemented with scikit-learn [24]. CatBoost was implemented with the Catboost [26] library, while ResNet and MLP were implemented with the tensorflow [30] module. The ResNet architecture was adapted from [31]. TabNet was implemented using PyTorch [32].

#### Influence of epitope length

In IFNBoost, we utilized a fixed length epitope sequence of 30 amino acids as input. Given that epitopes typically vary in length (see Supplementary Figure 1), we investigated whether incorporating epitope length as an additional input parameter would enhance model performance. However, our analysis indicated that epitope length has a minimal impact on the model’s predictions (Figure 4b left panel). Moreover, the input feature importance analysis revealed that epitope length is less significant than both the epitope sequence and the other three metadata features (Figure 4b right panel).

#### Robustness to amino acid encoding

In the IFNBoost model, we employed ordinal encoding of amino acids to represent epitope sequences. Ordinal encoding is a simple and efficient method, but many other encoding schemes can be used to represent epitope sequences, such as DPC [8] and those based on different physicochemical and structural properties of amino acids [22, 23]. Therefore, we tested whether the model’s performance could be improved by using these alternative encoding schemes. The different amino acid encodings utilized are DPC, physicochemical properties based encoding [22], and amino acid classification given by [23] (see **Materials and Methods** for details). Additionally, DPC was combined with the epitope length (DPC+length) as an additional feature to evaluate its impact on model performance.

IFNBoost demonstrated consistent performance across the various amino acid encodings (Figure 4c), suggesting that IFNBoost model is not overly reliant on any particular encoding scheme. The “DPC+length” encoding produced slightly better results compared to other encodings, indicating a marginal advantage when incorporating dipeptide composition along with length information. The improvement however is minimal, highlighting that amino acid encoding has a relatively limited impact on the model’s predictive performance. These observations reinforce the importance of epitope metadata over amino acid encoding in predicting IFN*γ* responses (Figures 4a). This also aligns with the observation that no differences in amino acid preferences were found between IFN*γ* inducing epitope sequences and non-inducers (Supplementary Figure 5). Also, since DPC encoding generates a vector of length 400 for each input sequence, compared to a length of 30 for other methods, it may lead to overfitting and reduced generalizability. Consequently, DPC also requires significantly more training time than the other methods.

### Comparative evaluation of computational models for predicting IFN*γ* inducing peptides

To ensure our choice of XGBoost is justified for IFNBoost, we assessed the performance of different computational models in predicting the IFN*γ* response. We assessed various approaches that include decision tree-based ML methods, neural network (NN) based methods, and Support Vector Machine (SVM). The decision tree-based methods included XGBoost [19], Random Forest [25], CatBoost [26] and AdaBoost [27], while the NN methods included residual neural network (ResNet) [28], deep neural network architecture for tabular data (TabNet) [29] and multi-layer perceptron (MLP). Similar to the IFNBoost model each method was fine-tuned using a five-fold CV on the training set containing 33,173 samples (see **Materials and Methods** for details). The performance of the models was evaluated on a common test dataset consisting of 8,294 samples (as in Figure 2b-2d).

Decision tree-based ensemble methods outperformed the NN based methods across all the evaluation metrics (Figure 5). All tree-based approaches achieved accuracy scores above 0.80, while all NN methods scored below 0.75. XGBoost (used in IFNBoost) exhibited the highest overall performance with an accuracy of 0.819. Among the compared methods, SVM performed the worst on our test set. Additionally, decision tree-based methods were faster to train compared to other methods.

### IFNBoost web server

We have developed a web server (https://ifnboost.streamlit.app/) for IFNBoost that facilitates text-based queries of epitopes (Supplementary Figure 6). It features a graphical user interface that takes 4 inputs from the user: epitope sequence, epitope species (source organism species), host, and assay method. The user needs to input a valid epitope sequence of length ranging from 12 to 30 amino acids. There is a drop-down menu to select among the three metadata features. The user can select from the available 43 hosts, 121 epitope species, and 7 assay methods, representing 99% of the samples in our training dataset.

The IFNBoost web server provides prediction results for a given epitope sequence, determining its likelihood of inducing an IFN*γ* response. The results include two components: prediction and prediction probability. The prediction provides the binary classification outcome (positive or negative IFN*γ* response) based on the model’s analysis. The prediction probability offers confidence levels for both outcomes in a tabular format. A higher prediction probability for a specific outcome indicates stronger confidence in the model’s output. This feature enables users to assess the reliability of the IFNBoost results and make informed decisions. Overall, this server serves as an easy-to-use platform for identifying IFN*γ* inducing epitopes, facilitating the development of rational vaccines and immunotherapies.

## Discussion

In this work, we introduced IFNBoost, an XGBoost based ML model designed for the rapid and accurate prediction of IFN*γ* inducing and non-inducing peptides, utilizing peptide sequence information and associated metadata. IFNBoost demonstrated excellent predictive performance on both the test and independent datasets, highlighting its accuracy and generalizability. An in-depth interpretation analysis of IFNBoost revealed the importance of different data factors in predicting IFN*γ* inducing peptides. IFNBoost is accessible as a web server for rapid deployment in the design of rational vaccines and immunotherapies.

IFNBoost uses three metadata features, host, source organism species, and assay method, that together enabled accurate prediction of IFN*γ* inducing peptides. These features are generally known to the user along with the epitope sequence information and do not require any additional experimental testing, making IFNBoost easily deployable for predicting IFN*γ* inducing peptides. Of the three metadata features, source organism species was found to be the most important feature (Figure 4a) as it seem-ingly narrows down the search space within a species during predictions.

IFNBoost outperformed the existing models for predicting IFN*γ* inducing peptides on an independent dataset across all evaluation metrics (Figure 3). This superior performance highlights the significance of incorporating metadata in identifying IFN*γ* inducing peptides, particularly as other methods rely exclusively on sequence information. It is widely recognized that the immune response against an epitope is influenced by several factors, including the host, source organism, method of observation, as noted in [33]. By incorporating these features in our model, we align with empirical evidence indicating that immune responses can vary for the same epitope sequence based on the metadata features. Notably, an epitope sequence may not induce IFN*γ* across different hosts, and even within the same host, an epitope can elicit differing IFN*γ* responses depending on the assay method employed (Supplementary Table 2). Incorporating epitope length as an additional feature in the model did not significantly enhance prediction accuracy (Figure 4b). This lack of improvement may be due to the ordinal encoding utilized in IFNBoost, which effectively captures epitope length through zero padding. Alternatively, it is possible that epitope length does not inherently contribute to distinguishing between the two classes, as suggested by our analysis of the epitope lengths of IFN*γ* inducers and non-inducers (Supplementary Figure 7).

The T cell epitope dataset used in this work comprises tabular data, organized into columns that provide detailed information about the epitopes. This type of data is inherently heterogeneous, with complex and irregular dependencies across columns. This is in contrast to image and natural language data which tend to have spatial dependencies and are more homogeneous [18]. Previous research has shown that decision tree-based methods often outperform deep learning NN methods on medium-sized (10,000 samples) tabular datasets [18, 34]. Our findings support this conclusion, as our comparison of computational models revealed that decision tree-based methods demonstrated superior performance compared to NNs. Additionally, decision tree-based methods offer the advantage of faster training times, a distinction we also observed during our analysis.

Gradient boosting methods, such as XGBoost, offer another advantage: they have an in-built mechanism to address missing data values by treating them as special cases during training to find optimal split directions, which are then consistently applied during predictions [19, 26]. This is valuable when certain metadata features are unknown, e.g., source organism species is not specified for some samples in our dataset. Thus, IFNBoost is adept at managing these anomalies, enhancing its applicability.

The IFNBoost model developed in this work for predicting IFN*γ* inducing peptides has several limitations. First, like any ML model, the performance of IFNBoost is dependent on the quality and quantity of available data. Second, limited data can result in models that are less generalizable and potentially biased. While IFNBoost performed well on an independent epitope dataset (Figure 3a), even for epitope sequences that were completely unseen by the model in the training data (Supplementary Figure 8), more extensive testing on unseen data will be required to further confirm its generalizability. Third, IFNBoost relies on multiple inputs, including three metadata features in addition to the epitope sequence. While this requirement may seem restrictive, this information is generally known to the user, and for cases where such metadata is unavailable, IFNBoost maps such inputs to an “others” category, ensuring the model remains functional and provides predictions based on the available information.

In this work, we specifically focused on identifying IFN*γ* inducing peptides due to their critical role in enhancing antigen presentation and promoting CTL response. Numerous other cytokines, such as IL-2, IL-5, IL-6, and TNF-*α*, are also known to play important, albeit distinct or complementary, roles in mounting immune responses against pathogens [1]. The generation of IL-6, for instance, is known to enhance antibody responses. Consequently, when designing effective antibody-based vaccines, it is essential to incorporate T cell epitopes that stimulate IL-6 production. Our proposed framework can be readily adapted to develop computational models for identifying peptides inducing specific cytokines, provided that there is a sufficient dataset available for both cytokine inducing and non-inducing peptides. This flexibility allows for a broader exploration of immune responses and the optimization of vaccine design.

The IFNBoost model can play a crucial role in the initial antigen selection phase of T-cell based vaccine design. By serving as a pre-screening tool, IFNBoost can identify peptides likely to induce IFN*γ* before synthesis and experimental testing, optimizing resource allocation. Successful peptides from experimental validation can rapidly advance to clinical trials, as preliminary computational predictions and subsequent ex-perimental validations provide robust evidence of their efficacy. IFNBoost can thus effectively guide the design of peptide-based T-cell vaccines for various diseases.

## Author contributions statement

A.A.Q. and M.S.S. conceived and designed the research. I.A. curated the datasets and developed the models. All authors contributed to data analysis and data visualization. A.A.Q. was responsible for funding acquisition and project supervision. I.A. wrote the original manuscript and A.A.Q. and M.S.S. reviewed and edited it. All authors discussed and approved the manuscript.

## Acknowledgments

We acknowledge all researchers at the originating and submitting laboratories that performed T cell assays and made the peptides available in IEDB. This work was supported by the Australian Research Council through the Discovery Project number DP230102850.

## References

[1] J. A. Owen et al., Kuby immunology, 7th ed., ser. Immunology. New York: W.H. Freeman, 2013.

[2] V. Schijns et al., “Rational vaccine design in times of emerging diseases: the critical choices of immunological correlates of protection, vaccine antigen and immunomodulation,” Pharmaceutics, vol. 13, no. 4, p. 501, Apr. 2021.

[3] R. Calderón-González et al., “Cellular vaccines in listeriosis: role of the Listeria antigen GAPDH,” Frontiers in Cellular and Infection Microbiology, vol. 4, p. 22, Feb. 2014.

[4] A. M. Gocher-Demske et al., “IFNγ-induction of TH1-like regulatory T cells controls antiviral responses,” Nature Immunology, vol. 24, no. 5, pp. 841–854, May 2023.

[5] G. Tau and P. Rothman, “Biologic functions of the IFN-γ receptors,” Allergy, vol. 54, no. 12, pp. 1233–1251, Dec. 1999.

[6] L. B. Ivashkiv, “IFNγ: signalling, epigenetics and roles in immunity, metabolism, disease and cancer immunotherapy,” Nature Reviews Immunology, vol. 18, no. 9, pp. 545–558, Sep. 2018.

[7] G. B. Klautau et al., “Interferon-γ release assay as a sensitive diagnostic tool of latent tuberculosis infection in patients with HIV: a cross-sectional study,” BMC Infectious Diseases, vol. 18, no. 1, p. 585, Nov. 2018.

[8] S. K. Dhanda et al., “Designing of interferon-gamma inducing MHC class-II binders,” Biology Direct, vol. 8, no. 1, p. 30, Dec. 2013.

[9] O. Singh et al., “ILeukin10Pred: A computational approach for predicting IL-10-inducing immunosuppressive peptides using combinations of amino acid global features,” Biology, vol. 11, no. 1, p. 5, Jan. 2022.

[10] M. T. Hassan et al., “Meta-IL4: An ensemble learning approach for IL-4-inducing peptide prediction,” Methods, vol. 217, pp. 49–56, Sep. 2023.

[11] A. Dhall et al., “A hybrid method for discovering interferon-gamma inducing peptides in human and mouse,” Scientific Reports, vol. 14, no. 1, p. 26859, Nov. 2024.

[12] R. Vita et al., “The Immune Epitope Database (IEDB): 2018 update,” Nucleic Acids Research, vol. 47, o. D1, pp. D339–D343, Jan. 2019.

[13] D. M. Tadros et al., “The MHC Motif Atlas: a database of MHC binding specificities and ligands,” Nucleic Acids Research, vol. 51, no. D1, pp. D428–D437, Jan. 2023.

[14] P. Wang et al., “A systematic assessment of MHC Class II peptide binding predictions and evaluation of a consensus approach,” PLOS Computational Biology, vol. 4, no. 4, p. e1000048, Apr. 2008.

[15] Y. EL-Manzalawy and V. Honavar, “Major histocompatibility complex (MHC), binder prediction,” in Encyclopedia of Systems Biology, W. Dubitzky et al., Eds. New York, NY: Springer, 2013, pp. 1162–1166.

[16] S. A. W. Shah et al., “Seasonal antigenic prediction of influenza A H3N2 using machine learning,” Nature Communications, vol. 15, no. 1, p. 3833, May 2024.

[17] R. Shwartz-Ziv and A. Armon, “Tabular data: Deep learning is not all you need,” Information Fusion, vol. 81, pp. 84–90, May 2022.

[18] V. Borisov et al., “Deep neural networks and tabular data: A survey,” IEEE transactions on neural networks and learning systems, vol. PP, Dec. 2022.

[19] T. Chen and C. Guestrin, “XGBoost: A scalable tree boosting system,” in Proceedings of the 22nd ACM SIGKDD International Conference on Knowledge Discovery and Data Mining, San Francisco California USA, Aug. 2016, pp. 785–794.

[20] T. Akiba et al., “Optuna: A next-generation hyperparameter optimization framework,” in The 25th ACM SIGKDD International Conference on Knowledge Discovery & Data Mining, 2019, pp. 2623–2631.

[21] J. Bergstra et al., “Algorithms for hyper-parameter optimization,” in Advances in Neural Information Processing Systems, vol. 24. Curran Associates, Inc., 2011.

[22] S. Kawashima et al., “AAindex: Amino acid index database,” Nucleic Acids Research, vol. 27, no. 1, pp. 368–369, Jan. 1999.

[23] M. J. Betts and R. B. Russell, “Amino-Acid properties and consequences of substitutions,” in Bioinformatics for Geneticists, Mar. 2007, pp. 311–342.

[24] F. Pedregosa et al., “Scikit-learn: Machine learning in Python,” Journal of Machine Learning Research, vol. 12, no. 85, pp. 2825–2830, 2011.

[25] L. Breiman, “Random Forests,” Machine Learning, vol. 45, no. 1, pp. 5–32, Oct. 2001.

[26] L. Prokhorenkova et al., “CatBoost: unbiased boosting with categorical features,” in Advances in Neural Information Processing Systems, S. Bengio et al., Eds., vol. 31. Curran Associates, Inc., 2018.

[27] Y. Freund and R. E. Schapire, “A desicion-theoretic generalization of on-line learning and an application to boosting,” in Computational Learning Theory, P. Vitányi, Ed., Berlin, Heidelberg, 1995, pp. 23–37.

[28] K. He et al., “Deep residual learning for image recognition,” in 2016 IEEE Conference on Computer Vision and Pattern Recognition (CVPR), Jun. 2016, pp. 770–778, iSSN: 1063-6919.

[29] S. Arik and T. Pfister, “TabNet: attentive interpretable tabular learning,” Proceedings of the AAAI Conference on Artificial Intelligence, vol. 35, pp. 6679–6687, May 2021.

[30] M. Abadi et al., “TensorFlow: A system for large-scale machine learning.”

[31] Y. Gorishniy et al., “Revisiting deep learning models for tabular data,” Advances in Neural Information Processing Systems, vol. 34, pp. 18932–18943, Dec. 2021.

[32] A. Paszke et al., “PyTorch: an imperative style, high-performance deep learning library,” in Proceedings of the 33rd International Conference on Neural Information Processing Systems. Red Hook, NY, USA: Curran Associates Inc., Dec. 2019, no. 721, pp. 8026–8037.

[33] J. Sidney et al., “Epitope prediction and identification-adaptive T cell responses in humans,” Seminars in immunology, vol. 50, p. 101418, Oct. 2020.

[34] L. Grinsztajn et al., “Why do tree-based models still outperform deep learning on typical tabular data?” Advances in Neural Information Processing Systems, vol. 35, pp. 507–520, Dec. 2022.

