## Supplementary Data for "IFNBoost: An interpretable computational model for identifying IFNγ inducing peptides"

### SUPPLEMENTARY FIGURES

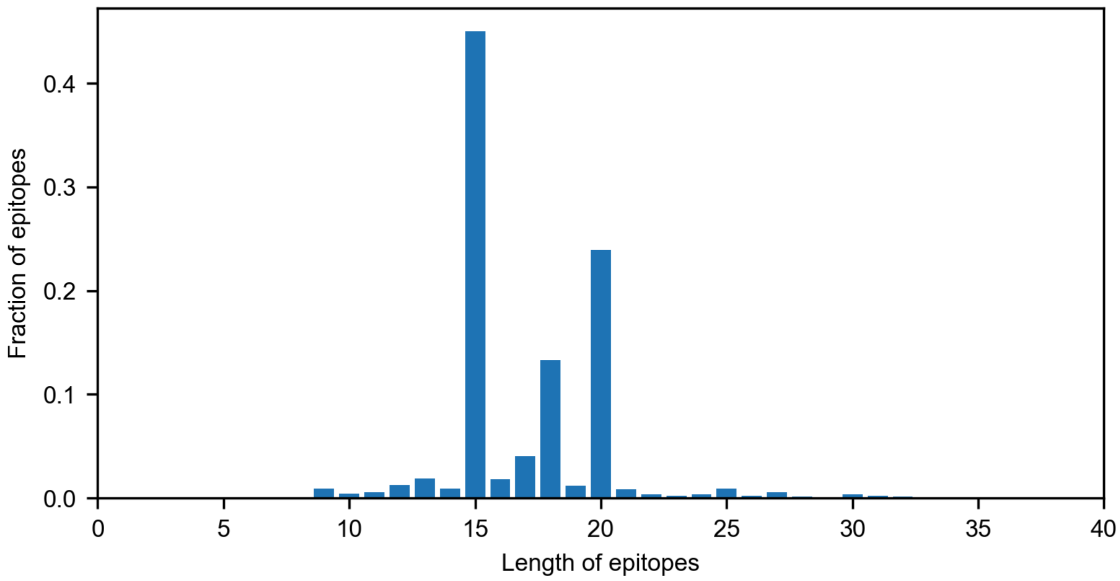


**Supplementary Figure 1: Distribution of epitope lengths in the dataset.** 45% of the epitopes have a length of 15 amino acids, indicating a significant preference for this length in the dataset. The majority of epitopes (~97%) lie within the range of 10 to 25 amino acids.

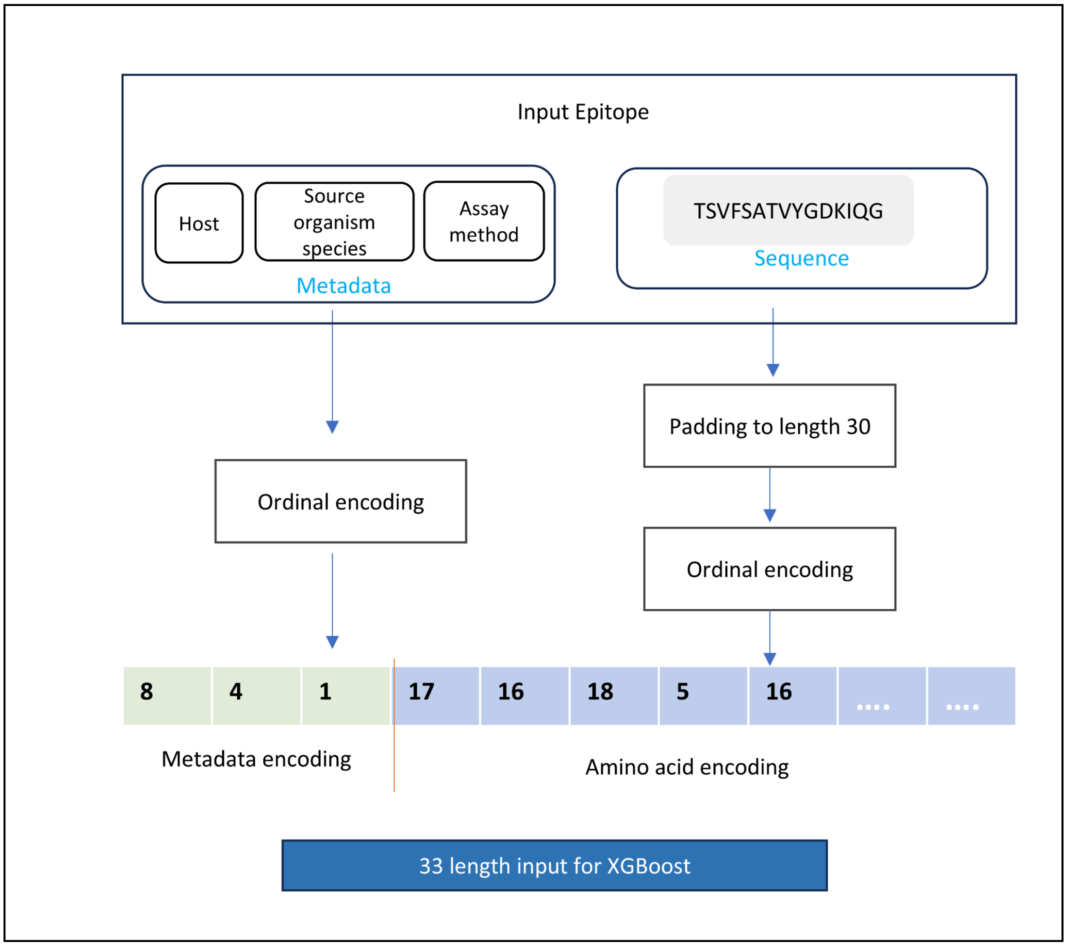


**Supplementary Figure 2: Feature encoding and input representation for the model**. Each input to the model is a vector of length 33, where the first three features are host, source organism species, and assay method. The remaining thirty represent the categorical encoding of amino acids. Both metadata and epitope sequence are encoded numerically to be provided as input to the XGBoost model.

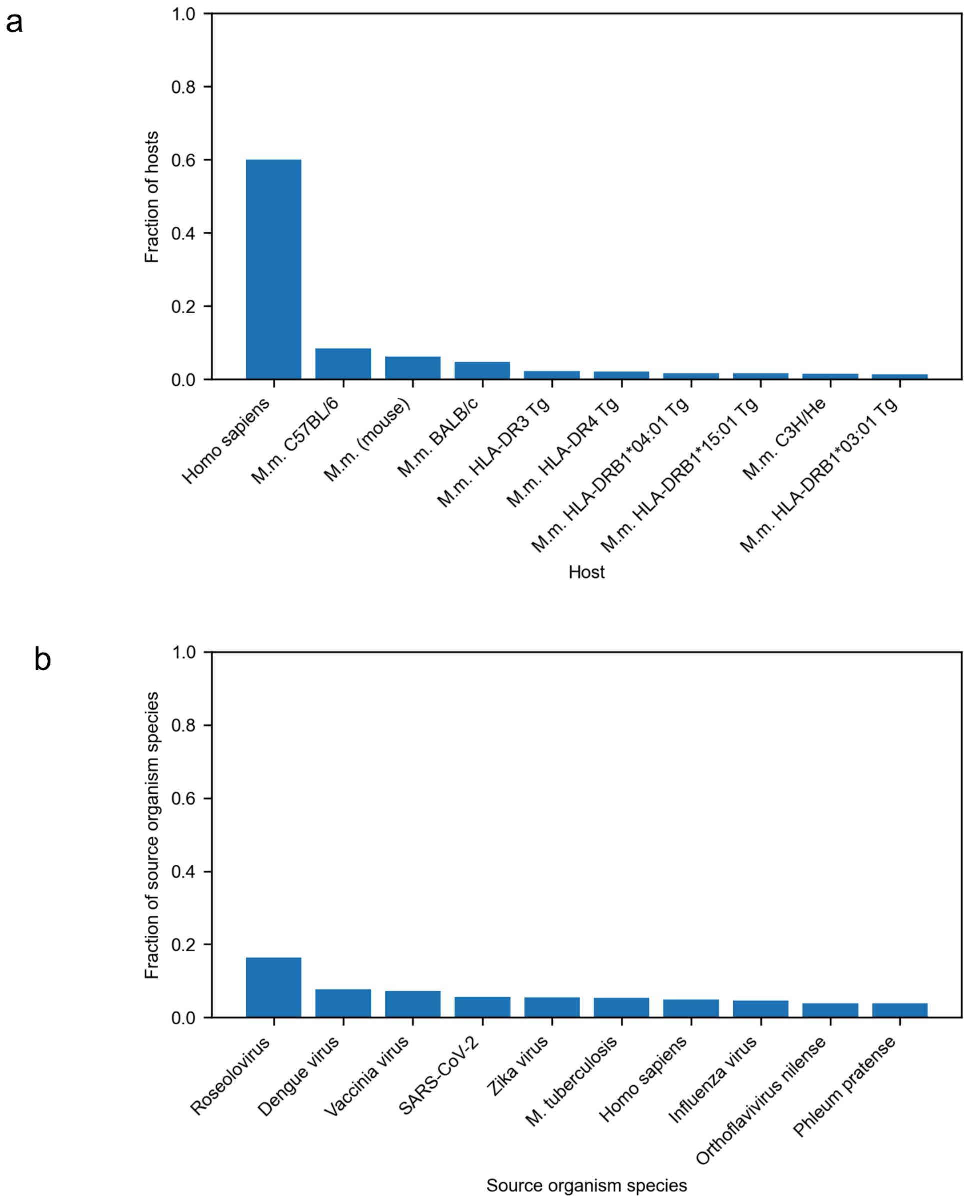


**Supplementary Figure 3: Data distribution of training set comprising 33,173 epitopes.** (a) The bar plot illustrates the fraction of top 10 hosts, which account for 89% of all epitopes in the training data. *Mus musculus* is represented by M.m. (b) The fraction of top 10 source organism species, which together represent 65% of the epitopes in the training data. *Mycobacterium tuberculosis* is abbreviated as *M. tuberculosis*.

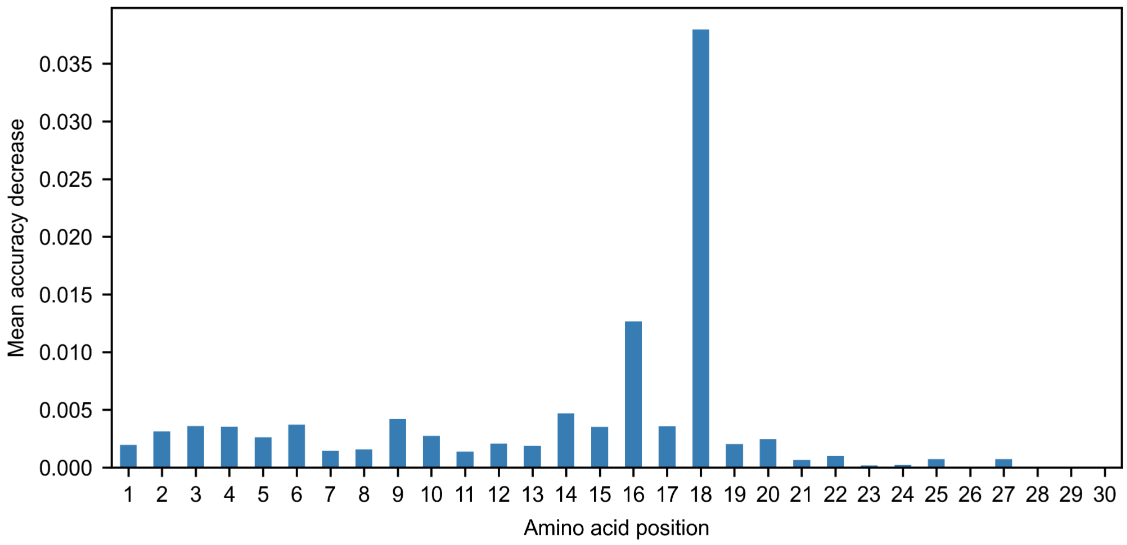


**Supplementary Figure 4: Input feature importance of IFNBoost.** The bar chart illustrates the permutation importance analysis of different amino acid positions in the input features of the model. The analysis was performed using the scikit-learn [1] library's ‘permutation_importance’ function, considering 33 input features. This plot presents the importance of 30 positions within the epitope sequence. Each bar represents the mean accuracy decrease by randomly shuffling a single feature value. The analysis involved 10 repetitions, i.e. the permutation process is repeated 10 times for each feature. Error bars are not included for clarity. The importance of IFNBoost has been calculated on the same test set used to report results in Figure 2b-2d.

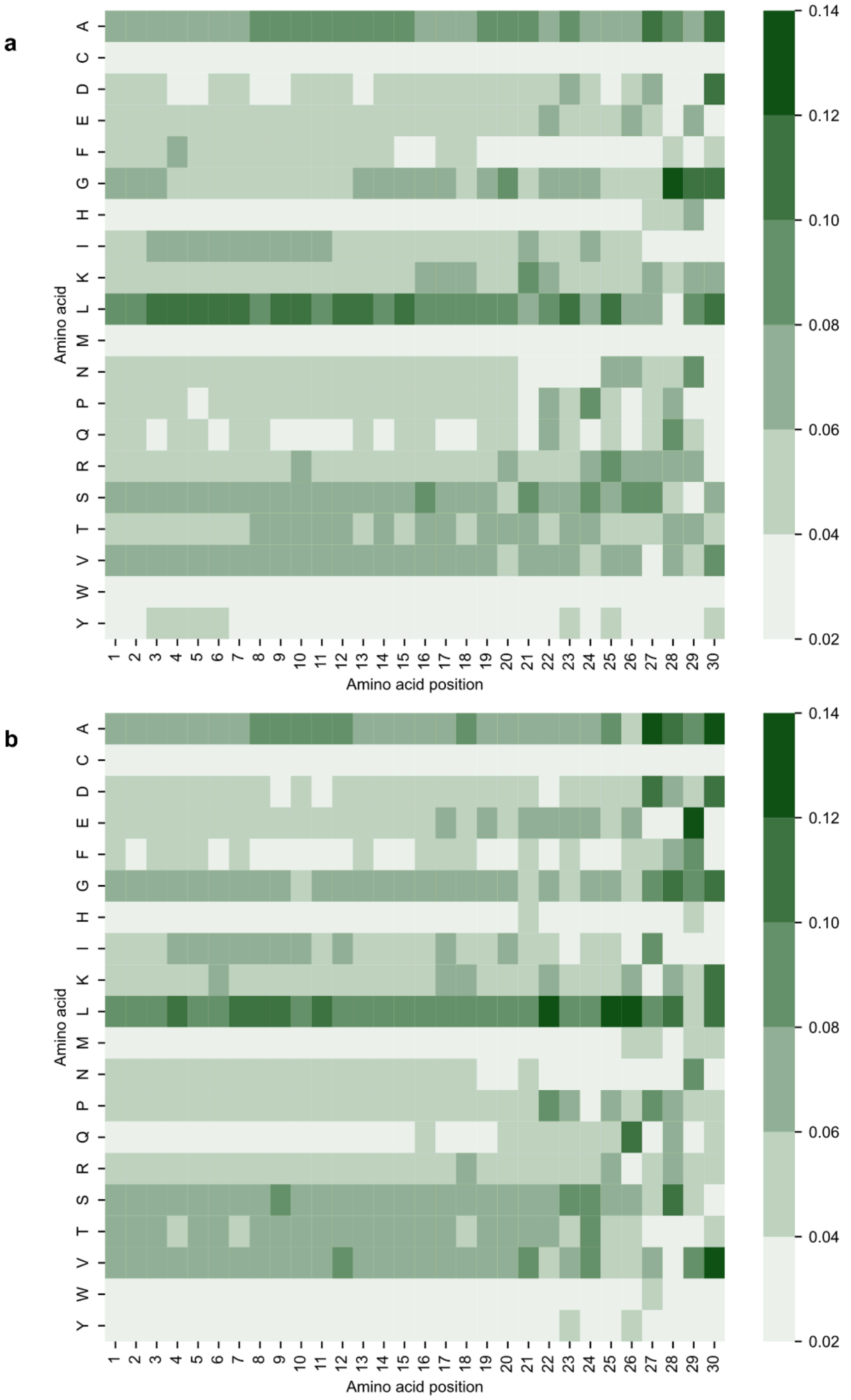


**Supplementary Figure 5: Amino acid preference at different epitope sequence positions. Statistics are shown for** a) IFNγ inducing and b) non-inducing epitopes in the dataset used for model development. Each cell in the heatmap corresponds to the fraction of a specific amino acid at a particular position within the epitopes. The color intensity reflects the fraction values, with a colormap ranging from low (light) to high (dark). No positional preference of amino acids is observed among the IFNγ inducers or non-inducers. L is the most frequent amino acid, followed by A at all positions. This implies that the amino acids at specific positions play a minor role in the identification of IFNγ inducing epitopes. We also computed the Bhattacharyya coefficient (BC) [2] to compare the probability distribution functions of amino acid counts at different positions within epitope sequences of IFNγ inducers and non-inducers. Our analysis yielded a BC of 0.95, indicating that there is a minimal distinction between the amino acid positions across the two classes.

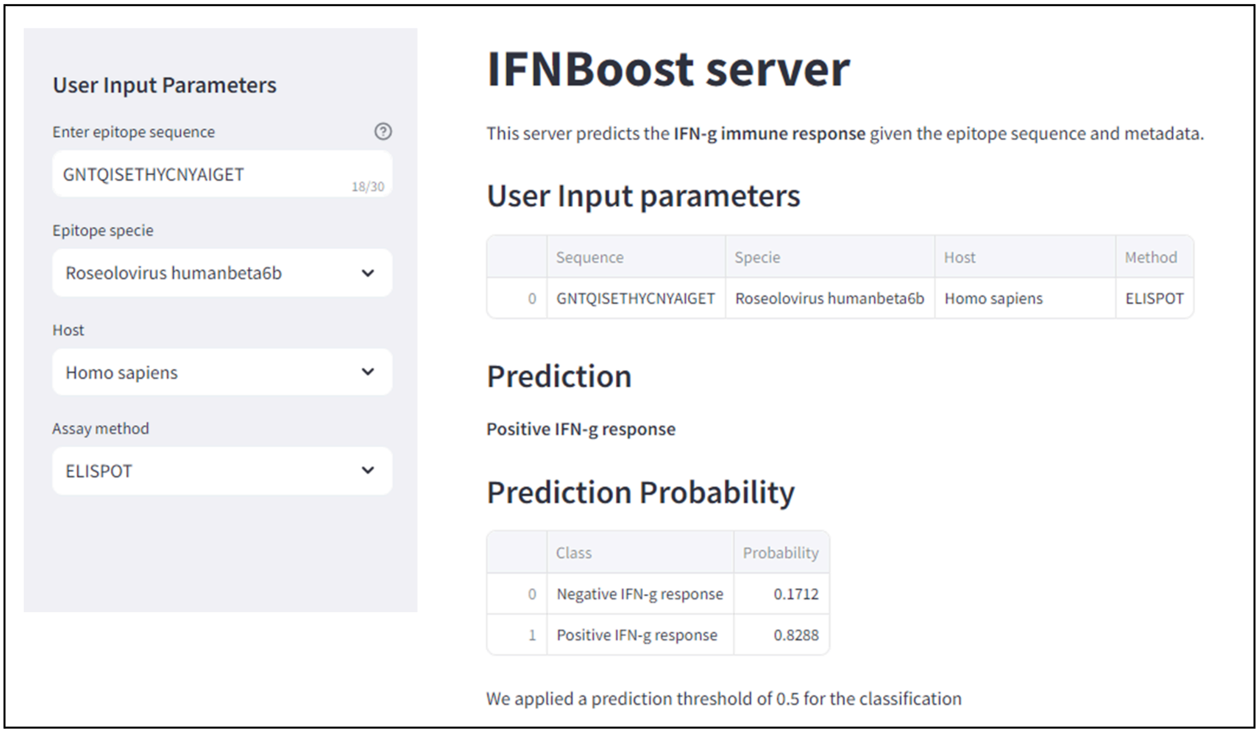


**Supplementary Figure 6**. **The IFNBoost web server user interface for predicting IFNγ inducing epitopes.** The left panel includes the four parameters that the user needs to input to run the IFNBoost model. An example case of a particular epitope is shown. The model prediction and the associated prediction probability is provided as outputs. The IFNBoost web server is publicly available at https://ifnboost.streamlit.app/.

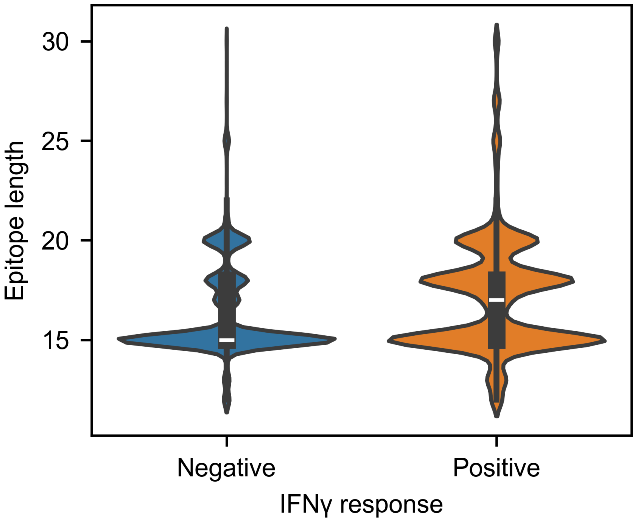


**Supplementary Figure 7: Epitope sequence length variations.** The violin plot represents the distribution of epitope lengths of IFNγ inducing epitopes and non-inducers respectively. The width of the violins indicates the density of epitope lengths, with wider regions signifying higher density. The length of both IFNγ inducers and non-inducers varies from 12 to 30 amino acids, with a predominant concentration (91.32%) observed in the range of 15 to 20 amino acids. We performed the Mann and Whitney U test [3] and found no statistically significant difference between the lengths of the IFNγ inducers and non-inducers (p-value = 0.3).

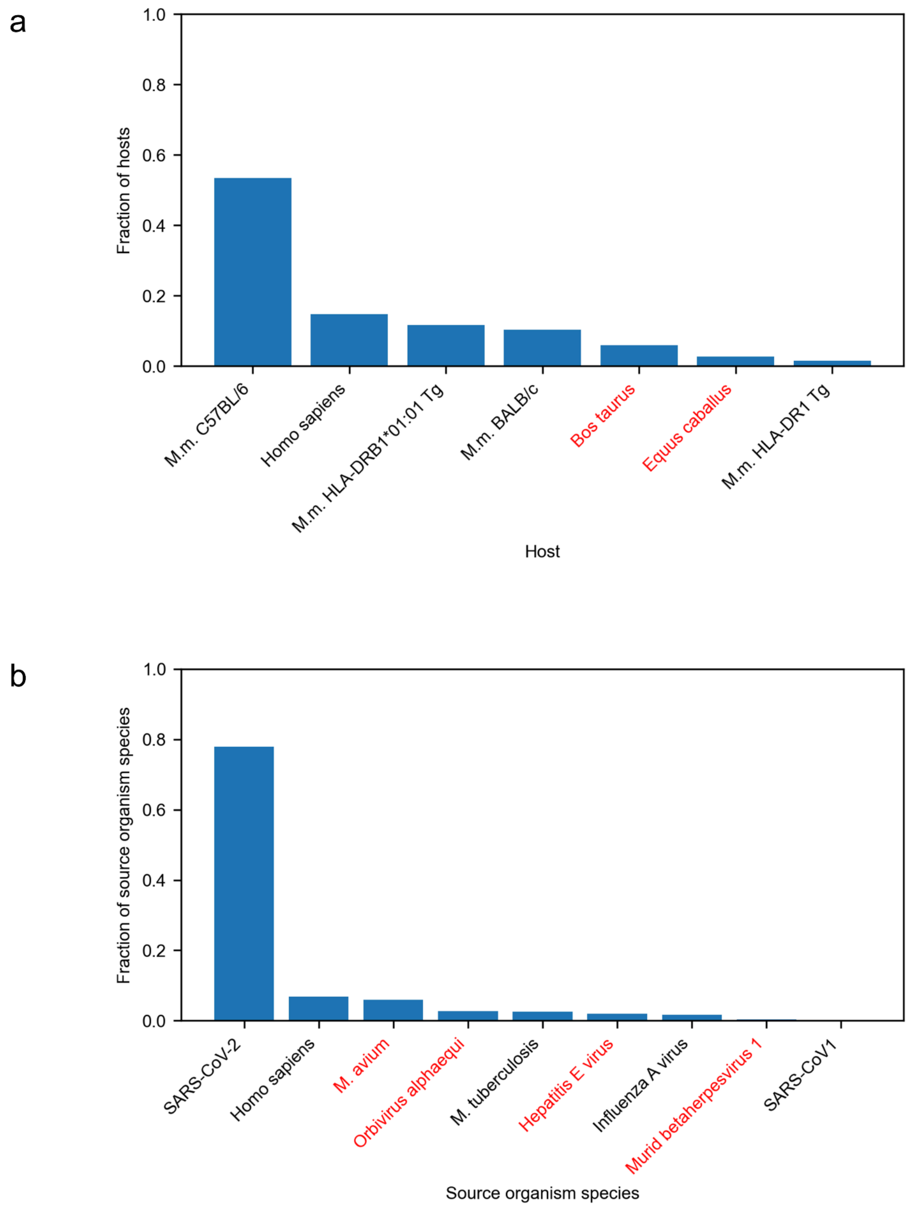


**Supplementary Figure 8: Statistics of the independent dataset from 2024.** This dataset comprised 1024 samples that were used to evaluate the performance of IFNBoost and existing IFNγ-inducing peptide prediction models in Figure 3. This dataset consisted of epitopes from seven hosts and nine source organism species. Entries highlighted in red denote novel categories that were not present in the training data of IFNBoost. (a) The fraction of epitopes from each of the seven hosts present in the dataset, with *Mus musculus* abbreviated as M.m.. (b) The fraction of epitopes from each of the nine source organism species, with *Mycobacterium* abbreviated as M..

### SUPPLEMENTARY TABLES

**Supplementary Table 1: Hyperparameter search space**

| Hyperparameter | Range | Increment |
|---|---|---|
| *n_estimators* | 100 - 1500 | 10 |
| *learning_rate* | 0.01 - 1 | 0.01 |
| *max_depth* | 1 - 100 | 1 |
| *lambda* | 0.1 - 10 | 0.1 |
| *alpha* | 0.1 - 10 | 0.1 |
| *gamma* | 0.1 - 10 | 0.1 |
| *min_child_weight* | 1 - 10 | 1 |
| *subsample* | 0.1 - 1 | 0.1 |
| *colsample_bytree* | 0.1 - 1 | 0.1 |

**Supplementary Table 2: Comparison of IFNγ response of same epitope sequence with different metadata features.**

| Epitope | Source organism species | Host | Assay method | IFNγ response |
|---|---|---|---|---|
| DRRWCFDGPRTNTIL | Orthoflavivirus nilense | **Mus musculus HLA-DRB1*04:01 Tg** | ELISPOT | Negative |
| DRRWCFDGPRTNTIL | Orthoflavivirus nilense | **Mus musculus HLA-DR3 Tg** | ELISPOT | Positive |
| ADLEVVTSTWVLVGGVLAAL | Hepacivirus hominis | Homo sapiens | **ELISPOT** | Positive |
| ADLEVVTSTWVLVGGVLAAL | Hepacivirus hominis | Homo sapiens | **ICS** | Negative |

### References

[1] F. Pedregosa *et al.*, “Scikit-learn: Machine learning in Python,” *J. Mach. Learn. Res.*, vol. 12, no. 85, pp. 2825–2830, 2011.

[2] A. Bhattacharyya, “On a measure of divergence between two multinomial populations,” *Sankhyā Indian J. Stat. 1933-1960*, vol. 7, no. 4, pp. 401–406, 1946.

[3] H. B. Mann and D. R. Whitney, “On a test of whether one of two random variables is stochastically larger than the other,” *Ann. Math. Stat.*, vol. 18, no. 1, pp. 50–60, 1947, doi: 10.1214/aoms/1177730491.
